# BCL6 is required for the development of functionally responsive IgM+ GC-independent Memory B Cells

**DOI:** 10.1101/564351

**Authors:** Gretchen Harms Pritchard, Akshay T. Krishnamurty, Lauren Rodda, Christopher Thouvenel, Derek J. Bangs, Courtney E. McDougal, David J. Rawlings, Marion Pepper

## Abstract

Humoral immunity depends upon long-lived, antibody-secreting plasma cells and memory B cells (MBCs). MBCs exhibit significant phenotypic and functional heterogeneity shaped by interactions with CD4+ T cells. It is currently unclear how specific CD4+ T cell interactions with B cells influence specific MBC subset generation. We used genetic ablation and antibody depletion to dissect key CD4+ T cell/B cell receptor-ligand pair interactions to define the critical signals that govern the development of specific MBC populations. While it has previously been suggested that germinal center (GC) experience is required for the development of CD73+CD80+ MBCs that rapidly form secondary plasmablasts, highly functional IgM+ MBCs differentiate in a BCL6 and CD4+ T cell-dependent, but GC-independent manner. BCL6 upregulation, in the presence or absence of a GC, can therefore serve as a predictor of long-lived, functional MBCs.

**One Sentence Summary:** Differential requirements of BCL6 and Tfh cells lead to the generation of functionally distinct memory B cell populations.

## INTRODUCTION

Humoral immunity is mediated by long-lived plasma cells (LLPCs) and memory B cells (MBCs) differentiated from B cells activated through B cell receptor (BCR) binding to cognate antigens (Inoue and Kurosaki, 2024; Plotkin, 2010). Pre-existing antibodies secreted by LLPCs can provide immediate defense against invading pathogens while MBCs maintain their BCR expression and rapidly differentiate into antibody-secreting plasmablasts (PBs) upon rechallenge (Akkaya et al., 2020). Phenotypically and functionally diverse MBC populations can form in response to immunization or infection in mice and humans (Callahan et al., 2024; Krishnamurty et al., 2016; Price et al., 2021; Seifert et al., 2015; Weisel et al., 2016; Yates et al., 2013; Zuccarino-Catania et al., 2014). We previously demonstrated in a murine model of infection that both IgG+ and IgM+ CD73+CD80+ MBCs rapidly form PBs in response to rechallenge, with CD73+CD80+ IgM+ MBCs (referred to as DP IgM+) responding faster than CD73+CD80+ IgG+ MBCs (referred to as DP IgG+ or DP swIg+) (Krishnamurty et al., 2016). A third population of expanded CD73-CD80- IgD+ MBCs, are more naïve-like and thought to predominantly contribute to a subsequent GC response that can repopulate the MBC pool (Callahan et al., 2024; Dogan et al., 2009; Krishnamurty et al., 2016; Mesin et al., 2020; Zuccarino-Catania et al., 2014). It is currently unclear how these distinct functionally responsive MBC populations form and how the transcription factor BCL6 may be involved.

The current paradigm of MBC formation suggests that activated B cells undergo iterative and complex interactions with CD4+ T cells that influence their fate. CD4+ T cells and B cells first engage at the T-B border and interfollicular regions of the secondary lymphoid organs. Some of these activated B cells subsequently form low-affinity GC-independent MBCs (Taylor et al., 2012b), whereas other activated B cells continue to interact with CD4+ Tfh cells in the GC (GC Tfh), where BCL6 is expressed by both B cell and Tfh cell populations (Vinuesa et al., 2016). Within the GC, class switch recombination (CSR), BCR somatic hypermutation (SHM), and testing by CD4+ T cells leads to affinity maturation and the downregulation of BCL6 and eventual exit of diversified class-switched MBC and LLPC populations (Papa and Vinuesa, 2018). The distinct signals that lead to the differentiation of specific MBC populations, particularly the DP IgM+ MBCs that have undergone SHM but not CSR, are unknown. Using various knockout mice and antibody-mediated temporal deletion, we tested how specific CD4+ T cell interactions and transcriptional programs can influence MBC fate. Our results demonstrate that both DP MBC populations require intrinsic expression of BCL6, but only the majority of the DP swIg+ MBCs require GC Tfh cells whereas an expanded population of DP IgM+ MBCs can form in the absence of GC Tfh cells.

## RESULTS and DISCUSSION

### B cell-intrinsic BCL6 expression is required for the development of *P.ch*-specific DP MBCs

As the hallmark transcription factor of the GC, BCL6 regulates many aspects of GC B cell program, such as localization, DNA repair, and inhibition of a plasma cell fate (Laidlaw and Cyster, 2021). To test the requirement of BCL6 for the development of DP MBC populations, we set up chimeras using congenically disparate WT and B cell conditional BCL6 deficient (BCL6BKO) bone marrow (see Methods) in order to create a system in which BCL6-sufficient and -deficient B cells could respond to infection in the same inflammatory environment. WT mice were infected with 10^6^ *P.ch* infected red blood cells (iRBCs) and the B cell response was examined at various time points thereafter. Previously published techniques were used to examine rare antigen-specific B cells specific for the carboxy terminus of the blood stage expressed merozoite surface protein 1 (MSP1) (Krishnamurty et al., 2016). After infection with *P.ch*, MSP1-specific BCL6BKO did not form GC B cells while MSP1-specific WT cells did (Fig. S1A), confirming the important role for BCL6 in GC B cell generation (Dent et al., 1997; Fukuda et al., 1997; Ye et al., 1997). The complete loss of GC B cells among the BCL6BKO population was reflected in a decrease of the total number of MSP1-specific B cells as well as MBCs (Fig. S1B).

Among the MBC populations at memory time points >40 days post-infection, BCL6 deficiency resulted in a loss of IgM+ MBCs and swIg+ MBCs (reduced by ∼50 and ∼98%, respectively), compared to WT MSP1-specific MBCs (Fig. 1A). Analysis of the expression of CD73 and CD80 on each subset further revealed a selective loss of the DP cells that accounted for a significant portion of the IgM+ and swIg+ MBCs that were lacking in the BCL6BKO population (both reduced by ∼99%) (Fig. 1B). We also tested to see if cre toxicity (Becher et al., 2018; Schmidt-Supprian and Rajewsky, 2007) was responsible for the defect that we saw in the transgenic B cells, rather than loss of BCL6, by infecting WT mice or Mb1cre+ mice with *P.ch.* We saw that the GC compartment was unaffected, neither were DP MBC populations (Fig. S1C,D). Intrinsic BCL6 expression is therefore critical for the formation of the GC and DP MBCs.

**Figure 1.**
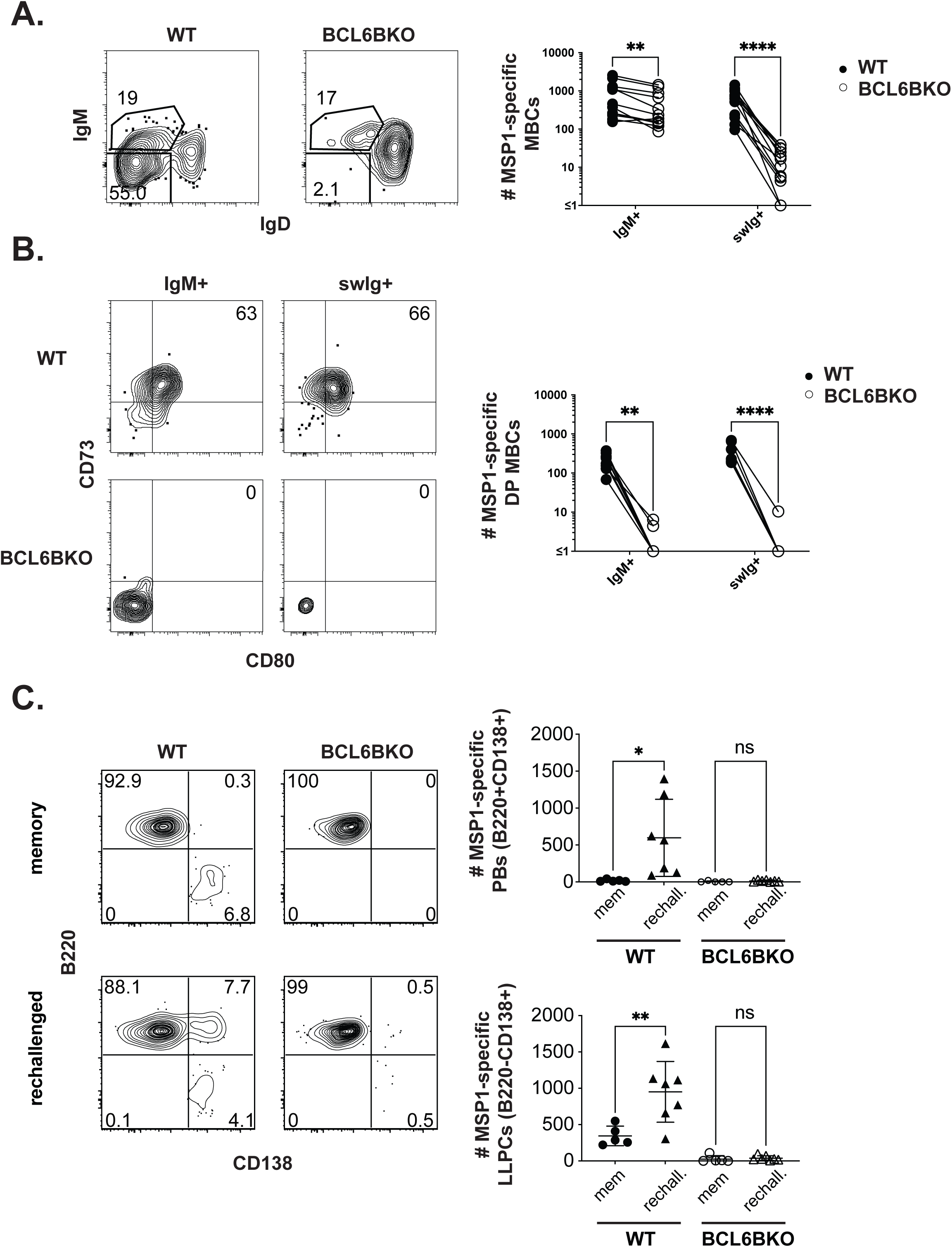
B cell intrinsic expression of BCL6 is required for IgM+ and swIg+ MBCs. Chimeras were made using congenically disparate WT (CD45.1/.2+) and BCL6BKO (CD45.2+) bone marrow mixed in equal proportions. After ≥8 weeks reconstitution, mice were infected with *P. chabaudi* and MSP1-specific B cell responses in the spleen were assessed at memory time points >40 days post-infection (A,B). MSP1-specific MBCs were assessed by isotype (A), and CD73 and CD80 expression (B). Data are pooled from 5 experiments with 1-5 mice per experiment. At memory time points (>40 days post-infection), mice were either unchallenged (memory) or rechallenged intravenously with 1x10^7^ iRBCs and MSP1-specific B cell responses in the spleen were analyzed by B220 and CD138 expression 3 days later (C). Data are pooled from 3 experiments with 1-3 mice per group in each experiment.

### BCL6-dependent DP MBCs are required for rapid PB formation

As previously mentioned, BCL6 controls many aspects of the GC B cell program including the negative regulation of several molecules associated with MBC phenotype including CD80 and PD-L2 (Niu et al., 2003; Peng et al., 2019). Therefore, while the preceding data demonstrated that BCL6BKO B cells do not form DP MBCs, it was possible that we had altered the phenotype but not the function of the MBCs. It was also possible that DP MBCs were not the only source of novel antibody secreting B220+CD138+ PBs (Krishnamurty et al., 2016). Thus, we compared functional responsiveness of WT and BCL6BKO cells to rechallenge. WT:BCL6BKO bone marrow chimeric mice were infected, allowed to form memory and subsequently rechallenged with *P.ch* at similar memory time points that we had previously seen functional PB responses to rechallenge (Krishnamurty et al., 2016). As expected, prior to rechallenge, WT B cells consisted of two populations of memory cells: B220+CD138- MBCs and B220^low^CD138+ LLPCs. LLPCs did not form in the BCL6BKO population as expected based on their GC-dependence (Fig. 1C) (Weisel et al., 2016). Similar to our previous results, WT cells formed an expanded B220+CD138+ PB population and an expanded LLPC population 3 days after rechallenge (Fig. 1C) (Krishnamurty et al., 2016). Yet both populations were absent in the BCL6BKO cells (Fig. 1C). B cell-intrinsic expression of BCL6 is therefore required for the formation of DP MBCs and LLPCs, and in the absence of these populations, no PBs are produced shortly after rechallenge with *P.ch*, again highlighting the importance of the DP MBC populations in a secondary response to infection.

### CD4+ T cells induce BCL6 expression in early activated B cells

While BCL6 is considered a “lineage defining” transcription factor of the GC, it is expressed during multiple phases of B cell activation. Elegant studies by Okada and colleagues using immunofluorescent microscopy of BCL6-YFP reporting B cells demonstrated that antigen-activated B cells initially up-regulate BCL6 in a T cell-dependent manner in the outer region of the follicle, before the GC has formed (Kitano et al., 2011). Moreover, studies from the Haberman laboratory identified perifollicular activated cells that express intermediate levels of BCL6 along with IRF4, and demonstrated that the cells that further upregulate BCL6 enter the GC whereas the cells that further upregulate IRF4 presumably form PBs (Zhang et al., 2017). We therefore sought to better understand the kinetics of BCL6 expression after infection, after excluding PBs from our analysis. WT mice were infected with 10^6^ *P.ch* infected red blood cells (iRBCs) and the infected-induced B cell response was examined at various time points thereafter.

MSP1-specific cells did not express BCL6 until day 6 post-infection, at which point IgD-BCL6+ cells began to accumulate (Fig. 2A). The majority of the IgD- BCL6-expressing cells were IgM+ six days post-infection yet the proportion of IgM+ BCL6+ cells decreased steadily and by day 20 post-infection BCL6+ cells were almost exclusively swIg+ (Fig. S2A). During this early activation period, CD73 and CD80 were both upregulated on IgD-BCL6- cells (Fig. 2B), suggesting the emergence of memory precursor cells (Kuo et al., 2007; Laidlaw et al., 2017).

**Figure 2.**
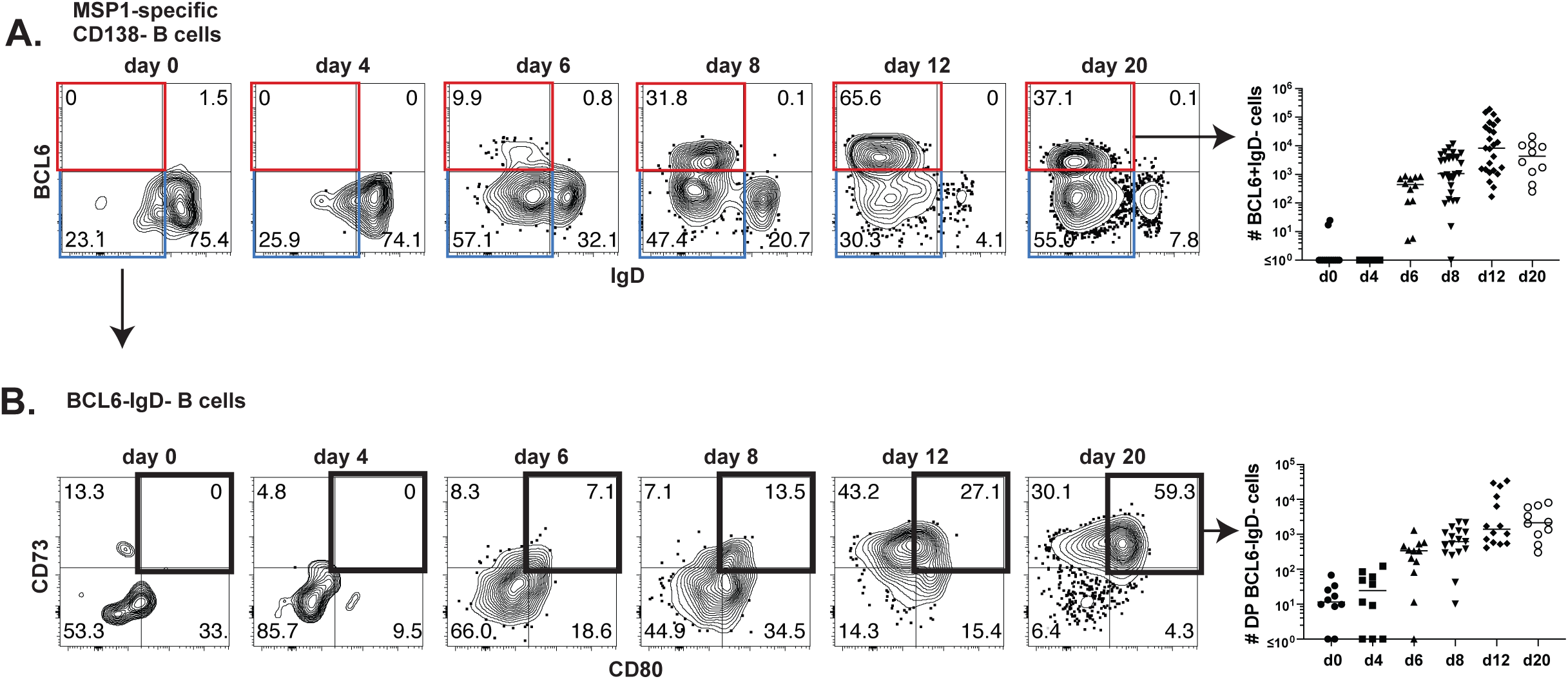
BCL6 is expressed in B cells after infection with *P. chabaudi*. WT mice were infected with *P. chabaudi* and MSP1-specific B cell responses were assessed at the indicated time points. Gating strategy and quantification of CD138-BCL6+IgD- cells (A). Gating strategy and quantification of CD73+CD80+ among BCL6-IgD- cells (B). Data are pooled from 9 independent experiments with 10-31 mice per time point.

We next asked whether early MSP1+ B cell BCL6 expression is dependent on CD4+ T cells, and if the early BCL6 expression drives the development of the subsequent DP MBC formation. To answer these questions, we first initiated treatment of mice with a CD4+ T cell-depleting antibody (clone GK1.5) one day prior to infection and examined the B cell response at day 7 post-infection. Depletion of CD4+ T cells prior to infection resulted in no MSP1 specific B cell BCL6 expression at day 7 whereas ∼25% of the CD138- B cells from the untreated controls did (Fig. 3A). Additionally, while CD4+ T cell depletion beginning on day 3 also impaired BCL6 expression, depletion at day 5 post-infection did not affect BCL6 expression at day 7 post-infection (Fig. 3A). Together with Figure 2A, these data demonstrate that BCL6 is upregulated by day 6 post-infection, and interactions with CD4+ T cells within the first 3 days of infection are essential for B cell upregulation of BCL6. Yet, when CD4+ T cells are depleted beginning at day 5 post-infection, BCL6 expression is maintained, indicating that delivery of the T cell signals inducing BCL6 upregulation have been initiated by this time point.

**Figure 3.**
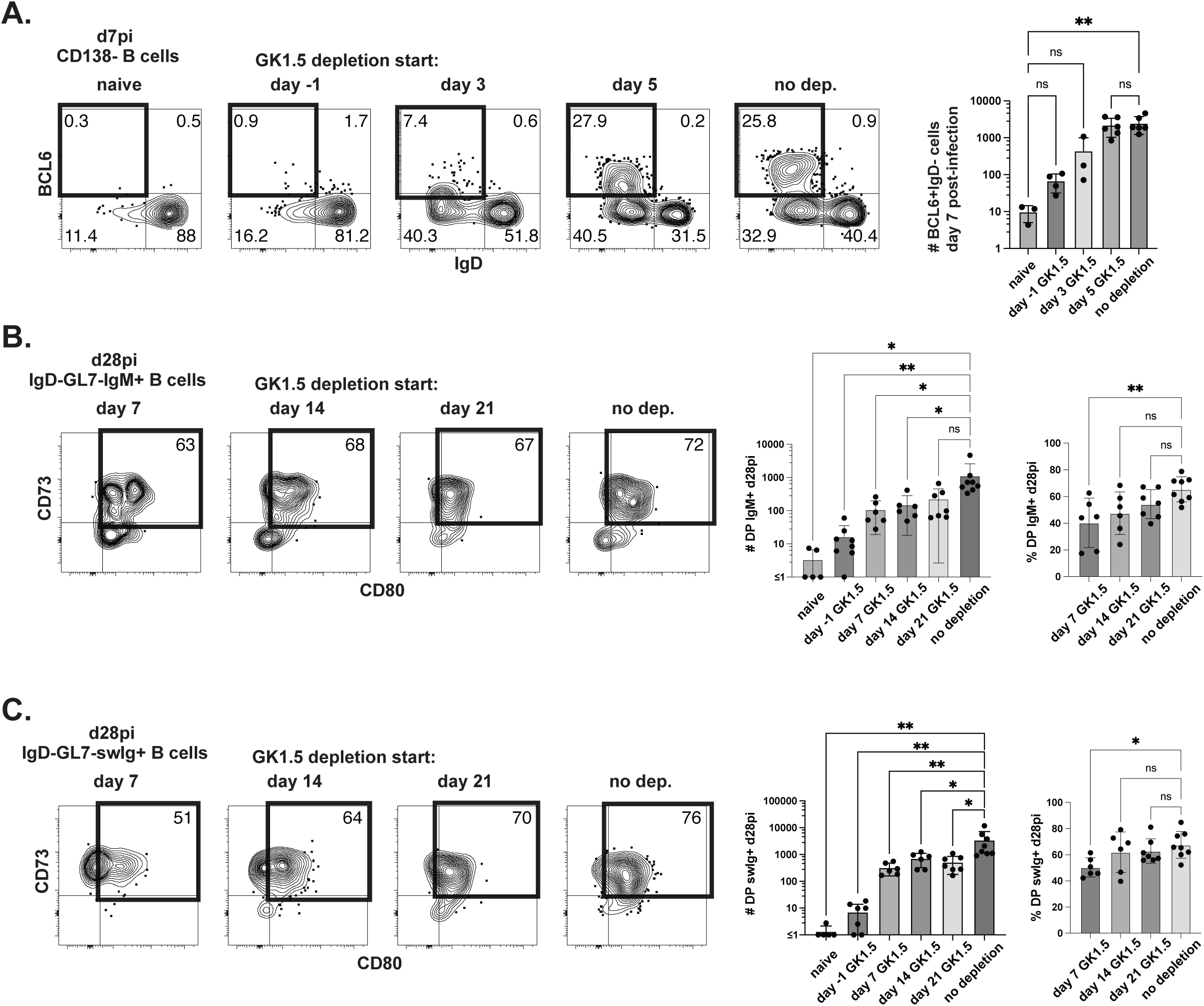
Early BCL6 expression is dependent on CD4+ T cells and required for DP MBC development. WT mice were treated with GK1.5 at the indicated time points during *P. chabaudi* infection and BCL6+IgD- cells were quantified at 7 days post-infection (A). Data are pooled from 2 independent experiments with 1-3 mice per group in each experiment. WT mice were treated with GK1.5 at the indicated time points during *P. chabaudi* infection and CD73 and CD80 expression was assessed on IgM+ (B) and swIg+ (C) GL7-IgD- cells. Data are pooled from 3 independent experiments with a total of 6-7 mice per time point.

In order to determine whether the CD4+ T cell dependent expression of BCL6+ occurring within the first week of infection was sufficient for the development of DP MBCs, or if CD4+ T cell interactions that occurred later than day 7 also contributed to the B cell response, we depleted CD4+ T cells at various time points of infection, and examined whether or not this led to distinct memory populations. Depletion of CD4+ T cells prior to infection resulted in very few IgM+ (mean=16.5, Fig. 3B) or swIg+ (mean=7, Fig. 3C) DP MBCs, reflecting a >98% reduction compared to untreated controls. These data demonstrate that CD4+ T cells are essential for the development of DP MBCs. While there was also a reduction in the number of DP MBCs in mice that had CD4+ T cells depleted beginning at all time points ≥ day 7 post-infection, the percentage of cells displaying CD73 and CD80 was largely intact, with no significant decrease in mice that received CD4+ T cell depleting antibody beginning at either day 14 or day 21, in both IgM+ (Fig. 3B) and swIg+ (Fig. 3C) MBC populations. Together, these data indicate that early after infection, CD4+ T cells deliver signals to B cells that initiate BCL6 upregulation and the subsequent development of DP MBCs, yet CD4+ T cell maintenance is required for the magnitude of the DP MBC populations.

### GC Tfh cells are not required for the development of somatically hypermutated DP IgM+ MBCs

Our data demonstrated that early interactions with T cells leads to the generation of BCL6-expressing B cells. We next asked what type of T cells are important for the development of DP MBC populations, specifically whether GC Tfh cells are required. Using a transgenic strain of *Plasmodium yoelii* that expresses the GP66 glycoprotein from LCMV (*P.y-GP66*), thus enabling us to track an antigen-specific T cell response, we have previously reported that by day 4 post-infection an expanded population of CXCR5 expressing T cells was detectible. By day 8 post-infection a subset of these cells had begun to upregulate PD-1, indicative of the emergence of a GC Tfh cell population (Arroyo and Pepper, 2020). Because B cell activation and upregulation of BCL6 followed similar kinetics to T cell activation and GC Tfh generation, we next sought to understand whether the early T cell signals were provided by GC Tfh cells. In order to test this, we generated mice with *T cell* intrinsic ablation of BCL6 (BCL6TKO) (see Methods), which is essential for the formation of GC Tfh cells (Hollister et al., 2013).

At 8 days post-infection with *P.ch*, there were equal numbers of CD38+GL7+ activated B cells between the WT and BCL6TKO mice (Fig. 4A). Furthermore, BCL6 was upregulated in B cells from both WT and BCL6TKO mice, although to a slightly diminished degree in the BCL6TKO cells (Fig. 4B). The data indicate that non- GC Tfh CD4+ T cells are able to provide the early signals leading B cell activation and BCL6 upregulation. To confirm that these early T cell signals can be delivered outside the context of a GC, we performed histology on spleens from WT and BCL6TKO mice at days 0 (i.e. naïve) and 6-8 post-infection with *P.ch* (Fig. 4C). In order to determine whether or not T cells had entered the B cell follicle we quantified size and T cell density of the early foci of BCL6+ cells surrounded by IgD+ follicular B cells. While both WT and BCL6TKO spleens contained early clusters within the follicle, they were smaller in the BCL6TKO mice and, as expected, contained a paucity of CD3+ cells (Fig. 4C). Together, these data demonstrate that while BCL6 expression within the T cell compartment is necessary for the development of a CD4+ GC Tfh cell population, B cells can become activated, express BCL6, and begin cluster coalescence in the absence of such cells.

**Figure 4.**
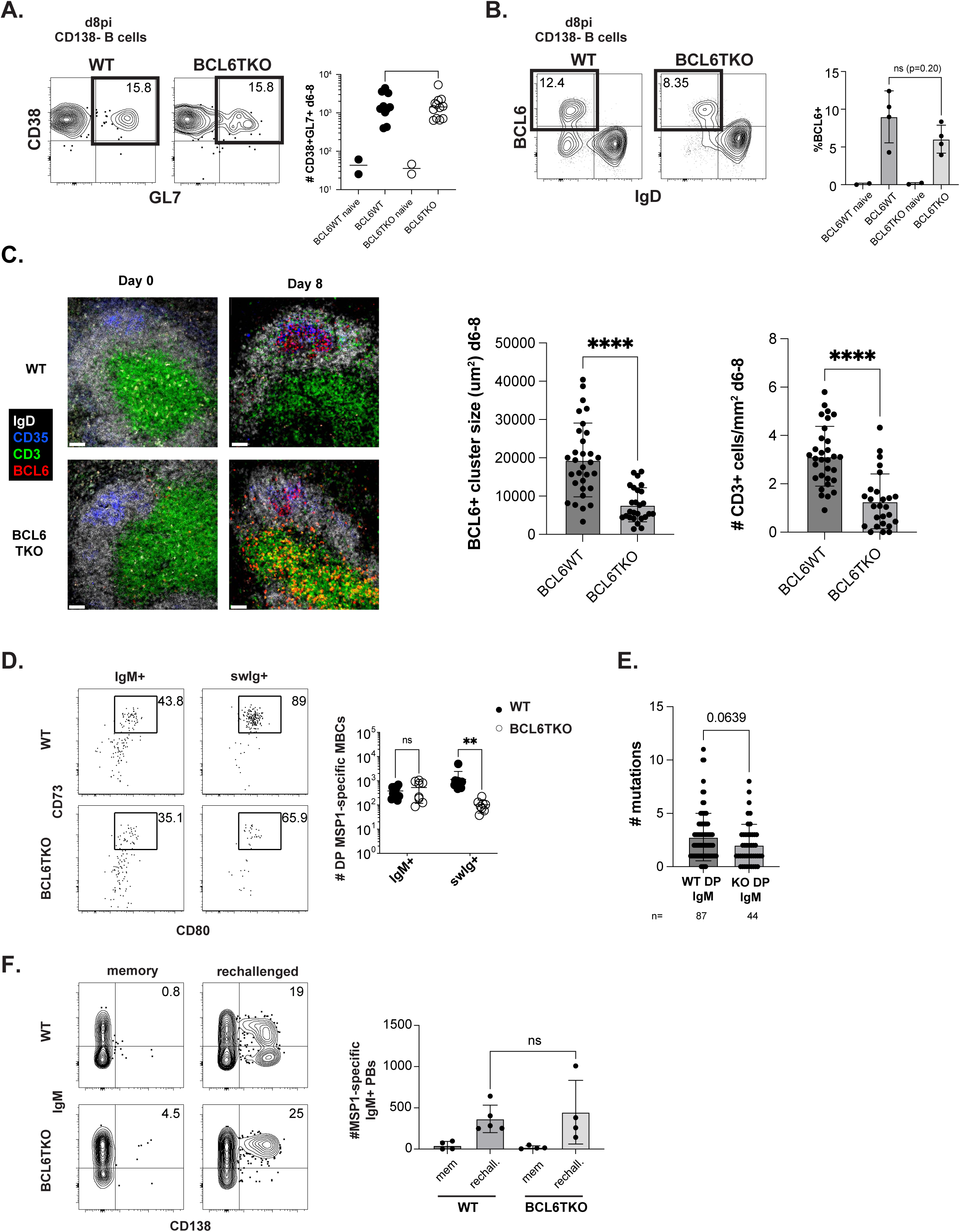
GC Tfh cells are not required for early B cell activation or the development of DP IgM+ MBCs. WT and BCL6TKO mice were infected with *P. chabaudi* and activation (CD38+GL7+ (A)) and BCL6 expression (B) was assessed at d8 post-infection. WT and BCL6TKO mice were infected with *P. chabaudi* and spleens prepared for histological analysis on 0 (n= 2 mice per genotype) and 6-8 (n = 3 mice per genotype) post-infection. The size and density of CD3+ cells in BCL6+ clusters from sections were quantified from d6-8 infected mice (C). Data are representative of 2 experiments and 3-30 follicles per mouse, 11-60 follicles per timepoint. Immunofluorescence microscopy of splenic B cell follicles. Scale bar is 50µm. WT and BCL6TKO mice were infected with *P. chabaudi.* At 28 days post-infection, CD73 and CD80 expression was assessed on IgM+ and swIg+ MBCs (D). Data are pooled from 3 individual experiments with 2-4 mice in each group. MSP1-specific DP IgM+ MBCs from 2 memory WT and 2 memory BCL6TKO mice were sorted and BCRs were sequenced and analyzed for the number of mutations among unique sequences (WT n=87, BCL6TKO n=44) compared to germline sequences (E). WT and BCL6TKO mice were infected with *P. chabaudi* and treated with atovaquone daily for 5 days starting at day 7 post infection. At ≥28 days post infection, mice were rechallenged with *P. chabaudi* and the IgM+ secondary plasmablast response was quantified 3 days later (F). Data are pooled from 2 independent experiments with 2-3 mice per group in each experiment.

Previous studies have described B cell clustering in the absence of T cells; however, these were in response to T-independent antigens and resulted in B cells that had not undergone SHM (de Vinuesa et al., 2000; Lentz and Manser, 2001). Therefore, we next analyzed the DP MBC response to determine if the B cells in the BCL6TKO mice were able to undergo SHM despite the absence of GC Tfh cells. By day 28 post-infection with *P.ch*, there were far fewer MSP1-specific B cells in BCL6TKO mice, and specifically, the GC B cell population was abolished as expected (Fig. S3A). Among MBCs, the DP IgM+ MBC population was intact yet there was a pronounced loss of DP swIg+ MBC in BCL6TKO mice compared to WT mice (Fig. 4D). We next performed BCR sequencing of sorted MBCs from *P.ch* infected WT and BCL6TKO mice and saw that BCRs from DP IgM+ MBCs from both WT and BCL6TKO had undergone SHM. However, there was a slight reduction in the level of SHM in the DP IgM+ MBCs from the BCL6TKO mice (mean number of mutations = 2.05) compared to the DP IgM+ MBCs from the WT mice (mean number of mutations = 2.78) (Fig. 4E). We were able to sort too few MSP1-specific DP swIg+ cells from the BCL6TKO mice (n=10) to make conclusions about this population (Fig. S3C). Together, these data demonstrate that the population of DP IgM+ MBCs can arise and undergo SHM in the absence of GC Tfh cells.

To address whether DP IgM+ MBCs can also develop independently of GC Tfh cells in other inflammatory settings or whether this phenotype was unique to *P.ch*, we examined the formation of PE-specific MBCs in mice immunized with PE protein in adjuvant. As seen with *P.ch* infection, the total number of PE-specific DP IgM+ MBCs was not affected by the loss of the GC Tfh compartment whereas the DP swIg+ MBCs were severely diminished (Fig. S3D). Together, these data demonstrate that while both DP IgM+ and DP swIg+ MBCs have undergone SHM, albeit to varying degrees, only the DP swIg+ MBCs require the presence of GC Tfh cells in order to expand. DP IgM+ MBCs can expand and undergo SHM outside of the GC reaction.

### The GC is not required for rapid secondary PB formation

Having established that the expanded DP IgM+, but not DP swIg+, MBCs can develop independently from the GC reaction, we next wanted to know whether these cells are able to form secondary PBs. Our previous experiments had compared the ability of different MBC populations to respond to secondary challenge within individual mice, however genetic ablation of the GC Tfh compartment necessitated comparing populations of cells in different groups of mice. When we assessed parasitemia in WT and BCL6TKO we found that there were no significant differences in parasitemia through day 30 post-infection, consistent with previous results (data not shown) (Pérez-Mazliah et al., 2017). However, to ensure that the PBs formed were induced by secondary infection rather than any sub-patent parasites that remained in the blood at undetectable levels, we treated WT and BCL6TKO mice with the antimalarial drug atovaquone prior to rechallenge. At 3 days post-secondary challenge we examined secondary PB formation. There was no significant difference between the ability of the WT and the BCL6TKO mice to mount a secondary IgM+ PB response (Fig. 4F), demonstrating the intact functionality of the GC Tfh-independent DP IgM+ MBCs. We also confirmed the ability of the GC Tfh deficient mice to mount a secondary PB response using the PE immunization setting and saw similar results (Fig. S3E).

Our data provide evidence that prior to entering the follicle and establishing GC formation, CD4+ T cells deliver activation signals that are sufficient to induce BCL6 upregulation. In the WT:BCL6BKO mixed chimeras, BCL6 deletion in the B cells results in ablation of both DP MBC populations. However, in mice without GC Tfh cells (i.e. BCL6TKO mice), the DP IgM+ MBC population is intact while the DP swIg+ MBC population is reduced, but not absent. Together, these data demonstrate that development of both of these populations is dependent on BCL6 and is induced prior to GC formation, yet the population of mutated DP MBCs that retain IgM expression bypasses the GC in its development.

The current model for the role of the GC in MBC development suggests that only low affinity, unmutated DN MBCs can be generated independently of the GC; in contrast, high affinity, mutated DP MBCs are dependent on the GC. Our approach of ablating BCL6 in either the B cell compartment or the T cell compartment enabled us to decouple the effects of intrinsic BCL6 expression and GC experience on the development of MBC populations. These data suggest that BCL6 has an essential function(s) in the development of MBC responses, independent of its role in GC formation. Indeed, BCL6 has myriad functions including regulation of B cell localization through repression of EBI-2, regulation of additional co-stimulatory molecule expression and protection from DNA-damage induced apoptosis (Kerfoot et al., 2011; Klein and Dalla-Favera, 2008; Pereira et al., 2009; Ranuncolo et al., 2007). Without this protection, B cells are not able to withstand DNA damage, such as that incurred during SHM and isotype class switch recombination (CSR). The GC is often considered to be the sole location of both of these functions (De Silva and Klein, 2015), yet in keeping with the work of others, our results demonstrate that both can take place outside of the GC. For example, Shlomchik and colleagues have reported that SHM can occur at the T zone-red pulp border (William et al., 2002), and somatically hypermutated swIg+ PBs are formed in extrafollicular sites during infection with *Salmonella typhirmurium* or *Ehrlichia muris* (Di Niro et al., 2015; Trivedi et al., 2019).

The studies presented here give us further insight into the intricacies of endogenous MBC development after infection and protein immunization and challenge current paradigms of MBC development.

## MATERIALS AND METHODS

### Mice

C57BL/6 (WT), B6.SJL-Ptprc^a^ Pepc^b^/BoyJ (CD45.1+), and B6.Cg-Tg(Cd4-cre)1Cwi/BfluJ (CD4-Cre+), mice were purchased from The Jackson Laboratory. CD45.1+ mice were crossed to WT mice to generate CD45.1+CD45.2+ mice. Mb1Cre+ mice were provided by Dr. Michael Reth (Max Planck Institute of Immunobiology and Epigenetics) and crossed to Bcl6flx/flx mice, provided by Dr. Alexander Dent (Indiana University), to generate Mb1-Cre+Bcl6flx/flx (BCL6BKO) mice. CD4-Cre+ mice were crossed to Bcl6flx/flx mice to generate CD4-Cre+ Bcl6flx/flx (BCL6TKO) mice. For experiments with BCL6TKO mice, Bcl6flx/flx Cre- littermates were used at WT controls. All mice were maintained/bred under specific pathogen-free conditions at the University of Washington. All experiments were performed in accordance with the University of Washington Institutional Care and Use Committee guidelines.

### Infections, drug treatment and parasitemia analysis

*Plasmodium chabaudi chabaudi (AS)* parasites were maintained as frozen blood stocks and passaged through donor mice. Primary mouse infections were initiated by intraperitoneal (i.p.) injection of 1x10^6^ iRBCs from donor mice. Secondary mouse infections were performed using a dose of 1x10^7^ iRBCs injected intravenously (i.v.). When indicated, mice were treated i.p. with 14.4mg/kg atovaquone resuspended in DMSO (Sigma) on days 7-11. When indicated, mice were i.p. injected with 250ug CD4 depleting antibody clone GK1.5 (BioXcell) beginning at the indicated time points and continued semi-weekly through the duration of the experiment. Parasitemia was measured by flow cytometry by fixing a drop of blood with 0.025% glutaraldehyde and staining with Ter119 FITC, CD45 APC, Hoechst33342, and CD71 PE. For memory PE experiments, mice were injected at the base of the tail with 15ug PE in CFA. For secondary PE responses, previously immunized mice were given 60ug PE in CFA i.p. (Pape et al., 2011).

### Bone Marrow Chimeras

Bone marrow cells were collected from the tibia, femur, humerus, and sternum and labeled with anti-Thy1.2 (30-H12, eBioscience) and anti-NK1.1 (PK136, eBioscience). Cells were resuspended and incubated with low toxicity rabbit complement (Cedarlane Laboratories). After complement lysis, cells were washed with media containing 10% fetal calf serum. Recipient mice were lethally irradiated (1000 rads) and injected intravenously with 5x10^6^ total bone marrow cells with congenically disparate WT cells mixed with BCL6BKO cells mixed in equal portions and provided with antibiotic (Enrofloxacin)-treated water for 8 weeks. Prior to infection, chimerism was determined analyzing the ratio of CD45.1 and CD45.2 expression among B220+ cells in the blood. In order to adjust for differences in initial chimerism, this ratio was then applied to the calculations obtained after infection.

### Tetramers

Purified recombinant His-tagged C-terminal MSP1 protein (amino acids 4960 to 5301) (Ndungu et al., 2009) was biotinylated and tetramerized with streptavidin-PE or streptavidin-APC (Prozyme), as previously described (Krishnamurty et al., 2016). Decoy reagent to exclude non-specific binding to the MSP1-PE tetramer was made by conjugating SA-PE to AF647 using an AF647 protein labeling kit (ThermoFisher), washing and removing any unbound AF647, and incubating with an excess of an irrelevant biotinylated His-tagged Class II-associated invariant chain peptide (Krishnamurty et al., 2016; Taylor et al., 2012a). Decoy reagent to exclude non-specific binding to the MSP1-APC tetramer was made by conjugating SA-APC to DyLight 755 using a DyLight 755 antibody labeling kit (ThermoFisher), washing and removing any unbound DyLight 755, and incubating with an excess of irrelevant biotinylated His-tagged Class II-associated invariant chain peptide.

### Mouse Cell Enrichment and Flow Cytometry

Splenic single-cell suspensions were prepared by mashing spleens and passing through 100um Nitex mesh (Amazon.com). For B cell enrichment, cells were resuspended in 200uL in PBS containing 2% FBS and Fc block (2.4G2) and first incubated with Decoy tetramer at a concentration of 10nM at room temperature for 20 min. MSP1-PE tetramer or MSP1-APC tetramer was added at a concentration of 10nM and incubated on ice for 30 min. Cells were washed, incubated with anti-PE or anti-APC magnetic beads for 30 min on ice, and enriched for using magnetized LS columns (Miltenyi Biotec). All bound cells were stained with surface antibodies followed by fixation and intracellular antibody staining when needed (Table 1). Cell counts were determined using Accucheck cell counting beads. All cells were run on the LSRII, Fortessa, Canto RUO or Symphony (BD) and analyzed using FlowJo software (Treestar).

### Bulk BCR Sequencing and Analysis

cDNA was generated from bulk sorted cells using a Template Switching RT Enzyme Mix (NEB) and the manufacturer’s protocol for Smart-seq at double volume per reaction. To account for larger cell numbers greater than 1000, a longer 17 base barcoded Template Switch Oligo (TSO) was used, and a shorter 8 base barcoded TSO was used for counts less than 1000. All reactions used a poly-T primer that shared the sequence of the TSOs and allowed for subsequent smart-seq amplification of all cDNA. A 1/3 portion of the barcoded cDNA was used in a qPCR reaction to determine the optimal number of cycles before plateau for each reaction. The rest of the barcoded cDNA was then amplified accordingly. To amplify BCRs, an Illumina sequencing method was used (manuscript in preparation as Thouvenel et al). Briefly, a multiplex PCR was performed on each bulk sample using pooled constant region primers for IgM, IgG, IgA, IgD, IgK, and IgL and sample-barcoded forward primers for the universal template switch region. Samples were then pooled and library prepped using enzymatic fragmentation and ligation with an anchored multiplex PCR approach (Zheng et al., 2014) to retain sample and transcript barcodes. Indexed library preps were run on an Illumina NovaSeq X at 300 cycles. Individual samples were demultiplexed by sample-barcoded sequences. UMI-tools (Smith et al., 2017) was used to create whitelists of transcript barcodes for each sample after manually determining the barcode counts from the knee in the rank abundance plots. TRUST4 (Song et al., 2021) was run for each sample along with the barcode whitelist to create BCR contigs. Contigs were submitted to IMGT/HighV-Quest (GENE-DB Version 3.1.42) for annotation and nucleotide mutation counts were parsed using an in-house python script from the output tables. Sequences were filtered by length so that all included at least the CDR1 to the end of the J-gene. Sequences for each sample were collapsed by V-gene, J-gene, nucleotide junction and amino acid mutations, and plotted in R (Version 4.4.1, http://www.r-project.org/) using the tidyverse (2.0.0, (Wickham et al., 2019) and ggplot2 (3.5.1).

### Histology

Spleens were fixed in 4% PFA for 2 hours at 4°C, washed three times for 10 min in 1X PBS and sunk in 30% sucrose overnight before freezing–in OCT. 7µm cryosections were dried for 1 hour and rehydrated in 1X PBS with 1% BSA for 10 min. Sections were stained in antibody overnight at 4°C, washed for 10min in 1X PBS with 1% BSA and stained in secondary antibody for 2 hours at room temperature. All stains were done in 1X PBS with 2% normal mouse serum (Jackson Immunoresearch), 2% normal rat serum (Jackson Immunoresearch) and 1% BSA. Antibodies used were anti-IgD ef450 (11-26c, eBioscience), anti-CD3 Alexa488 (17A2, BioLegend), anti-CD35 biotin (8C12, BD Pharmingen), anti-BCL6 Alexa647 (K112-91, BD Biosciences) and streptavidin Cy3 (Jackson Immunoresearch). Immunofluoresence microscopy images were captured with a Nikon Eclipse Ti microscope and analyzed with NIS-Elements AR.

### Statistical Analysis

Two-tailed Student’s *t* tests were applied to determine the statistical significance of the differences between individual groups. One-way ANOVA or two-way ANOVA was used to determine the statistical significance of the differences between multiple groups. All analyses were done with Prism (Graphpad) software. The p-values were considered significant when p < 0.05 (*), p < 0.01 (**), and p<0.001 (***).

## Supporting information

Antibody list

## List of Supplementary Materials

**Figure S1. Heterogeneous B cell populations are dependent on intrinsic expression of BCL6.** WT:BCL6BKO chimeric mice were infected with *P. chabaudi* and MSP1-specific GC B cells (A) and MBCs (B) were quantified at days 12 and >40 post-infection, respectively. WT and Mb1Cre+ mice were infected with *P. chabaudi* and MSP1-specific GC B cells (C) and DP MBCs (D) were analyzed ≥28 days later.

**Figure S2. BCL6 is expressed in both IgM+ and swIg+ B cells.** Isotype was assessed among BCL6+IgD- cells at the indicated time points (A).

**Figure S3. GC B cells do not develop in BCL6TKO mice and are not required for the DP IgM+ MBC response after PE immunization.** WT and BCL6TKO mice were infected with *P. chabaudi* and total MSP1-specific (A) and GC (B) B cells were quantified at day 28 post-infection. At 28 days post-infection with *P. chabaudi* DP IgG+ MBCs were sorted, BCRs were sequenced (WT n=141, BCL6TKO n=10), and mutation counts were determined by comparing to germline (C). WT and BCL6TKO mice were immunized with PE and PE-specific MBC isotypes were analyzed for CD73 and CD80 expression (D). WT and BCL6TKO mice were immunized with PE and allowed to form memory, then rechallenged with PE. PE-specific plasmablast responses were quantified 3 days later (E).

Table1. List of antibodies used

## Acknowledgements

The authors would like to thank Brian Hondowicz and Brian Johnson for technical assistance. We would also like to thank Dr. Michael Reth (Max Planck Institute of Immunobiology and Epigenetics) for providing the Mb1Cre+ mice, and Dr. Alexander Dent (Indiana University) for providing the Bcl6flx/flx mice. The authors have no conflicting financial interests.

## Author contributions

GHP conceptualized, performed and analyzed experiments, prepared figures, wrote and edited the manuscript, ATK conceptualized, performed and analyzed experiments, LR performed and analyzed experiments and prepared figures, CT and DJB analyzed experiments and prepared figures, CEM performed experiments, DJR contributed to experimental design and analyses, MP conceptualized and analyzed data from experiments and wrote and edited the manuscript.

