## Supplementary figures and images for "BCL6 is required for the development of functionally responsive IgM+ GC-independent Memory B Cells"

### Supplemental Figure 1

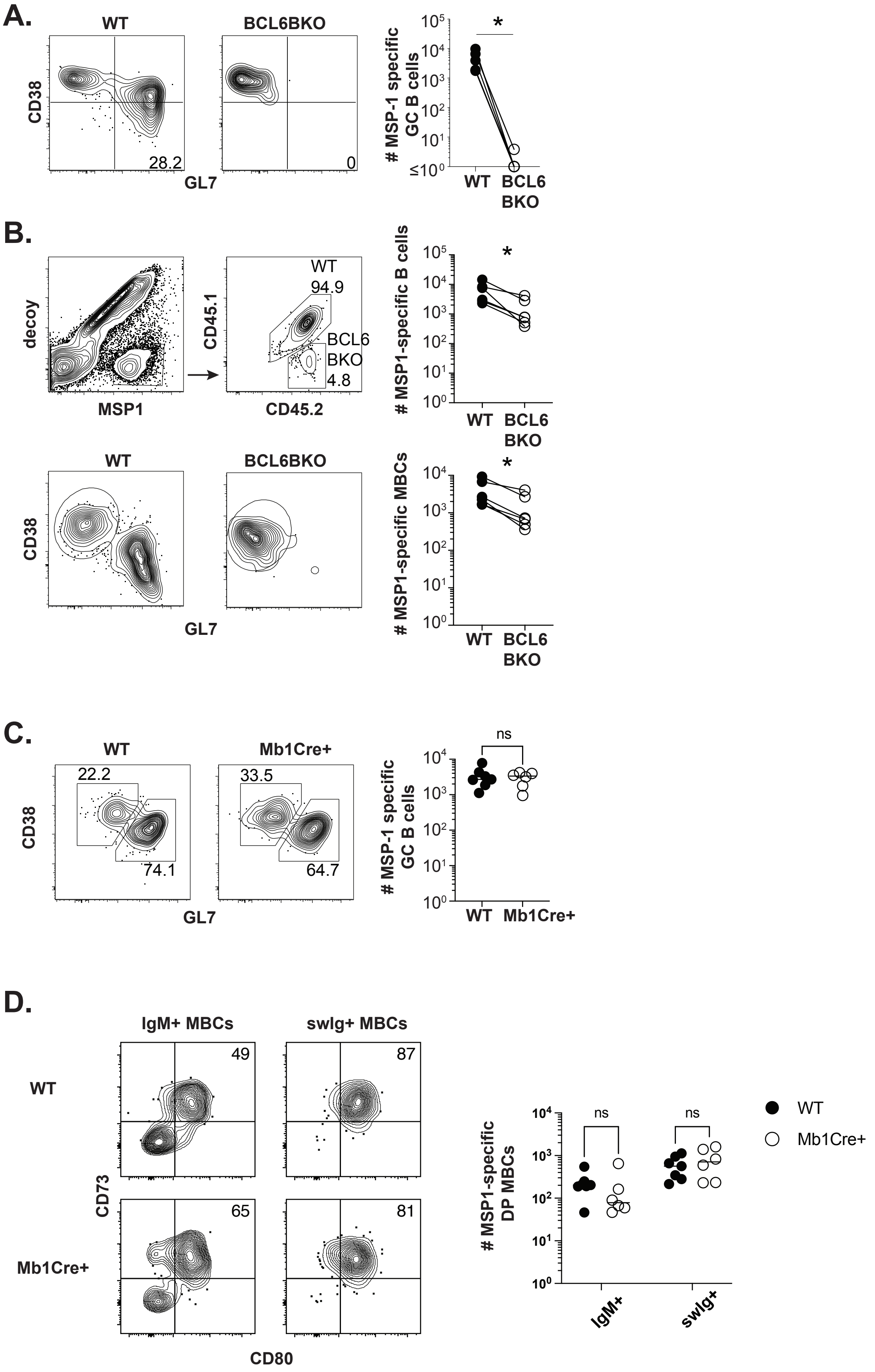

### Supplemental Figure 2

#### Fig. S2

## A.

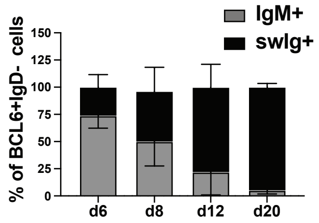

### Supplemental Figure 3

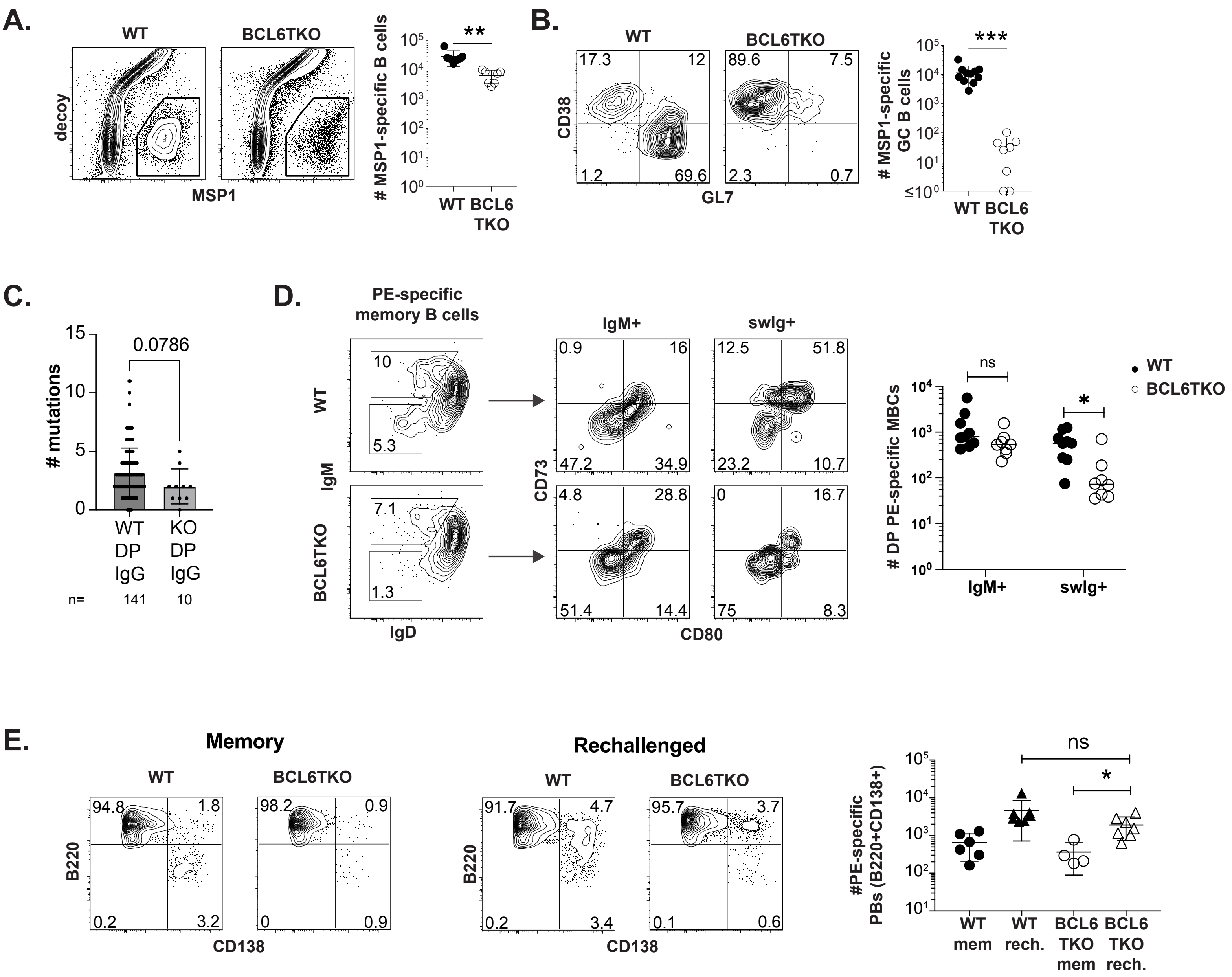
