## Supplementary material for "BCL6 is required for the development of functionally responsive IgM+ GC-independent Memory B Cells": Antibody list

Table1.

Antibodies used:

| Specificity | Conjugate | Clone | Company |
| --- | --- | --- | --- |
| B220 | BV711 | RA3-6B2 | BD Horizon |
| B220 | AF647 | RA3-6B2 | eBioscience |
| B220 | BUV737 | RA3-6B2 | BD Horizon |
| B220 | BV510 | RA3-6B2 | BD Horizon |
| BCL6 | AF647 | K112-91 | BD Pharmingen |
| CD138 | BV605 | 281-2 | BD Horizon |
| CD138 | BV650 | 281-2 | BD Horizon |
| CD138 | BB515 | 281-2 | BD Horizon |
| CD3 | PerCP-Cy5.5 | 145-2C11 | BD Pharmingen |
| CD3 | FITC | 145-2C11 | BD Pharmingen |
| CD38 | AF700 | 90 | Invitrogen |
| CD4 | BV510 | RM4-5 | BD Horizon |
| CD4 | biotin | GK1.5 | Pharmingen |
| CD4 | BV711 | GK1.5 | BD Horizon |
| CD4 | BUV805 | GK1.5 | BD Horizon |
| CD45.1 | APC-ef780 | A20 | eBioscience |
| CD45.2 | FITC | 104 | BD Pharmingen |
| CD45.2 | BV510 | 104 | BioLegend |
| CD45.2 | APC | 104 | BD Pharmingen |
| CD35 | biotin | 8C12 | BD Pharmingen |
| CD73 | PE-Cy7 | eBioTY/11.8 | eBioscience |
| CD73 | ef450 | eBioTY/11.8 | eBioscience |
| CD8 | AF647 | 53-6.7 | eBioscience |
| CD8 | ef450 | 53-6.7 | Invitrogen |
| CD8 | BV510 | 53-6.7 | BD Horizon |
| CD80 | FITC | 16-10A1 | BD Pharmingen |
| CD80 | BV605 | 16-10A1 | BD Horizon |
| CD80 | Biotin | 16-10A1 | BD Pharmingen |
| GL7 | ef450 | GL-7 | Invitrogen |
| GL7 | biotin | GL-7 | eBioscience |
| IgD | BV786 | 11-26c.2a | BD Horizon |
| IgD | BV650 | 11-26c.2a | BioLegend |
| IgD | BUV395 | 11-26c.2a | BD Horizon |
| IgM | APC | 11/41 | BD Pharmingen |
| IgM | BV786 | 11/41 | BD OptiBuild |
| SA | Cy3 |  | Jackson ImmunoResearch |
| SA | BV510 |  | BDVector |
| SA | APC-Cy7 |  | BD Pharmingen |
| SA | AF594 |  | Jackson ImmunoResearch |
| SA | BUV395 |  | BD Horizon |
